# The molecular genetics of participation in the Avon Longitudinal Study of Parents and Children

**DOI:** 10.1101/206698

**Authors:** Amy E. Taylor, Hannah J. Jones, Hannah Sallis, Jack Euesden, Evie Stergiakouli, Neil M. Davies, Stanley Zammit, Debbie A. Lawlor, Marcus R. Munafò, George Davey Smith, Kate Tilling

## Abstract

**Background:** It is often assumed that selection (including participation and dropout) does not represent an important source of bias in genetic studies. However, there is little evidence to date on the effect of genetic factors on participation.

**Methods:** Using data on mothers (N=7,486) and children (N=7,508) from the Avon Longitudinal Study of Parents and Children, we 1) examined the association of polygenic risk scores for a range of socio-demographic, lifestyle characteristics and health conditions related to continued participation, 2) investigated whether associations of polygenic scores with body mass index (BMI; derived from self-reported weight and height) and self-reported smoking differed in the largest sample with genetic data and a sub-sample who participated in a recent follow-up and 3) determined the proportion of variation in participation explained by common genetic variants using genome-wide data.

**Results:** We found evidence that polygenic scores for higher education, agreeableness and openness were associated with higher participation and polygenic scores for smoking initiation, higher BMI, neuroticism, schizophrenia, ADHD and depression were associated with lower participation. Associations between the polygenic score for education and self-reported smoking differed between the largest sample with genetic data (OR for ever smoking per SD increase in polygenic score:0.85, 95% CI:0.81,0.89) and sub-sample (OR:0.95, 95% CI:0.88,1.02). In genome-wide analysis, single nucleotide polymorphism based heritability explained 17-31% of variability in participation.

**Conclusions:** Genetic association studies, including Mendelian randomization, can be biased by selection, including loss to follow-up. Genetic risk for dropout should be considered in all analyses of studies with selective participation.

## Key messages

- Polygenic scores for a range of sociodemographic, health and lifestyle factors are related to continued participation after enrolment in the Avon Longitudinal Study of Parents and Children.
- There was evidence that associations between polygenic scores and measured phenotypes differed between the full sample with genetic data and a more selected sub-sample, indicating that genetic association studies can be biased by selection.
- Common genetic variation explained a moderate amount (17-31%) of variability in participation.
- Researchers should consider selective participation as a potential source of bias in genetic association studies.

## Introduction

Missing data are a pervasive problem in cohort studies, with decreasing participation over the duration of the study, and concern about the extent to which this biases analyses (1, 2). Individual characteristics, including social and lifestyle characteristics may influence both initial enrolment and continued participation (3, 4). Throughout this paper we use the word “participation” to mean both initial enrolment in a study and continued participation (e.g. via questionnaire completion or attendance at research clinics) once involved. However, our analyses all relate to continued participation after enrolment.

Sample representativeness is critical for estimating prevalence of exposure or disease (5), but may not be essential for estimating associations between exposures and outcomes (5–7). The bias arising from selection into studies is often relatively small and may not always qualitatively affect interpretation of results (1, 8, 9). Selection bias might be less problematic in genetic epidemiology because individuals are generally unaware of their genotype (so will not self-select into a study on the basis of this) and genetic variants that influence a given trait should not be associated with confounding factors which could also influence selection (6, 10). However, when both exposure and outcome relate to participation in a study, this can induce spurious associations between them, or between genetic variants that influence them, in participants (11, 12). For example, the association between higher genetic risk for schizophrenia and reduced participation in the Avon Longitudinal Study of Parents and Children (ALSPAC) (13) indicates that selection bias may be a problem in both genetic and non-genetic analyses of schizophrenia.

To estimate the impact of selective participation for a given analysis, we need to know which factors cause participation. Here, we extend previous work relating participation and polygenic risk for schizophrenia and autism in ALSPAC (13, 14) by 1) investigating polygenic scores for other factors which could influence participation in the ALSPAC mothers and children,2) investigating the potential impact of selection bias by comparing associations between genetic factors and measured phenotypes in the largest sample with genetic data and a more selected sub-sample and 3) conducting genome-wide association studies of participation measures.

## Methods

### Study population

ALSPAC is a longitudinal birth cohort that recruited 14,541 pregnant women resident in Avon, UK with expected dates of delivery from 1st April 1991 to 31st December 1992. Of these initial pregnancies, there were a total of 14,676 foetuses, resulting in 14,062 live births and 13,988 children who were alive at 1 year of age. The children and their mothers have been followed up through postal questionnaires and at clinics (3, 15). We included only children who had been enrolled in the study during the first phase of data collection and survived to age 1 year (resulting in the exclusion of 5 children and 43 mothers from the analysis sample). Please note that the study website contains details of all the data that is available through a fully searchable data dictionary: http://www.bris.ac.uk/alspac/researchers/data-access/data-dictionary. Ethical approval for the study was obtained from the ALSPAC Ethics and Law Committee and the Local Research Ethics Committees

### Participation

Participation was defined by responding to a questionnaire or attending a clinic for which the whole cohort was eligible to participate (i.e. we excluded clinics and questionnaires targeted at a subset of the cohort). The ALSPAC mothers have answered questionnaires about themselves (mother questionnaires) and about their children (child-based questionnaires). The ALSPAC children have answered questionnaires about themselves (child-completed questionnaires). A full list of the questionnaires and clinics included is provided in Supplementary Table S1. From these, we calculated the following continuous phenotypes by summing the number of questionnaires/clinics completed: total participation (all questionnaires and clinics for both mother and child (including child-based and child-completed)), total questionnaire (all questionnaires for mothers and children), mother questionnaire (mother questionnaires), child questionnaire (child-completed questionnaires), and child clinic (child clinics attended). We created two binary variables for the mothers and children indicating 1) participation in the most recent clinic and 2) completion of the most recent questionnaire. For both mothers and the offspring we generated variables from data collected at clinics 17-18 years after the child’s birth and from questionnaires 19-20 years after birth.

### Genetic data

ALSPAC children were genotyped using the Illumina HumanHap550 quad chip genotyping platforms. ALSPAC mothers were genotyped using the Illumina human660W-quad array at Centre National de Genotypage (CNG) and genotypes were called with Illumina GenomeStudio. Imputation was performed using Impute V2.2.2 against the 1000 genomes phase 1 version 3 reference panel. Quality control procedures removed related individuals and individuals of non-European genetic ancestry (see supplementary materials for full details).

### Polygenic scores

We calculated polygenic scores for a number of traits that could be related to participation and for which genome-wide summary statistics were publicly available: body mass index (16), height (17), smoking initiation (18), depression (19), attention deficit hyperactivity disorder (ADHD)(20), bipolar disorder (21), autism (21), schizophrenia (22), years of education (23), sleep duration (24), chronotype (morningness) (24), age at menarche (25), personality traits (openness, agreeableness, conscientiousness, extraversion and neuroticism) (26) and Alzheimer’s disease (27). For the purposes of this paper, we use the term “trait” to describe the phenotype each GWAS was conducted on, but acknowledge that for binary phenotypes, we are looking at genetic liability for that phenotype. Full details of sources for each of these scores are shown in supplementary Table S2. The ALSPAC cohort was not included in the GWAS that generated the summary statistics for these traits, except for education and age at menarche. For education, we used summary statistics excluding ALSPAC and 23andme, which were obtained directly from the study authors. For age at menarche, the ALSPAC sample made up 7% of the GWAS discovery sample (25). To minimise potential bias from sample overlap, we used an unweighted polygenic score for age at menarche (28). All other scores were weighted according to the association magnitude of each SNP in the original GWAS.

### Statistical analysis

All analyses were performed separately in mothers and children.

### Polygenic scores

Polygenic scores were derived using the PRSice software (http://prsice.info/) (29) for each trait within the ALSPAC genome-wide data using the following p-value thresholds: 0.0005, 0.005, 0.05, 0.1, 0.5 (see Supplementary Methods). In addition, we generated scores in PRSice by inputting only the independent genome-wide significant SNPs reported by the discovery samples (Supplementary Table S3). We assessed associations of standardized polygenic scores with participation phenotypes using linear and logistic regression in Stata (version 14.1)(30). We used robust standard errors to account for the non-normal distribution of the continuous participation variables. For age at menarche, analyses were conducted in females only.

### Genome-wide association analysis

Analyses were conducted separately for mothers and children. We used SNPTEST (31) to test associations between dosage scores for each genetic variant and missingness phenotypes using univariate regression models and assuming an additive genetic model. Continuous phenotypes were initially tested in linear models, and then dichotomised at arbitrary midpoints (Supplementary Table S4) and re-tested in logistic models to ensure results were robust to any assumption on the distribution of residuals. Genome-wide results were filtered to remove SNPs with a minor allele frequency of <0.01 and imputation quality (info) score of <0.8. Genome-wide significance was considered to be p<5x10^−8^ (32).

### Heritability

SNP-based heritability estimates *h^2^_SNP_* were calculated for each participation phenotype using the genetic restricted maximum likelihood (GREML) method implemented within the GCTA software (33).

### Investigating the impact of selection bias in ALSPAC

We used linear and logistic regression to calculate associations between polygenic scores for BMI, smoking, education and schizophrenia (constructed at a p-value threshold of 0.05) and body mass index and smoking status (ever vs never smoking) which were self-reported by the ALSPAC mothers in questionnaires administered during pregnancy. These analyses were conducted first in the largest sample with genome-wide data and then in the sample attending the most recent clinic.

## Results

Of the 13,793 mothers with 13,988 children alive at one year, 11,560 mothers and 10,780 children had provided DNA samples. After removal of non-Europeans, related individuals and samples which did not pass quality control, 7,486 mothers and 7,508 children were eligible for analysis (Table 1, Supplementary figures S1 and S2). Individuals included in the analysis had higher participation levels than the enrolled cohort (Supplementary Table S5). Continuous participation phenotypes were highly correlated (Pearson’s correlation coefficients ranged between 0.71 and 0.99) (Supplementary Table S6).

**Table 1.**
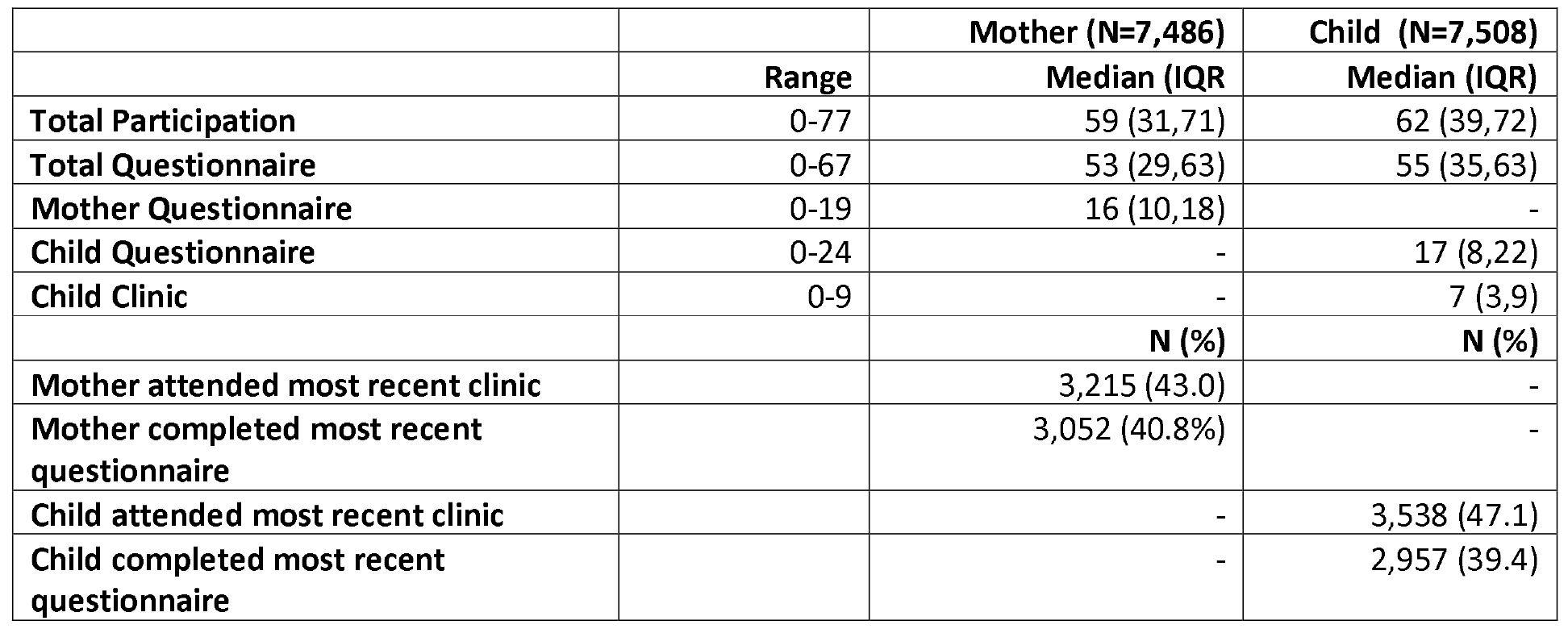
**Summary of participation phenotypes**

|  |  | **Mother (N=7,486)** | **Child (N=7,508)** |
| --- | --- | --- | --- |
|  | **Range** | **Median (IQR)** | **Median (IQR)** |
| **Total Participation** | 0-77 | 59 (31,71) | 62 (39,72) |
| **Total Questionnaire** | 0-67 | 53 (29,63) | 55 (35,63) |
| **Mother Questionnaire** | 0-19 | 16 (10,18) | - |
| **Child Questionnaire** | 0-24 | - | 17 (8,22) |
| **Child Clinic** | 0-9 | - | 7 (3,9) |
|  |  | **N (%)** | **N (%)** |
| **Mother attended most recent clinic** |  | 3,215 (43.0) | - |
| **Mother completed most recent questionnaire** |  | 3,052 (40.8%) | - |
| **Child attended most recent clinic** |  | - | 3,538 (47.1) |
| **Child completed most recent questionnaire** |  | - | 2,957 (39.4) |

### Associations of polygenic scores with participation phenotypes

Only the results for total participation and last questionnaire completion are presented, with results for all other participation measures in supplementary material.

In ALSPAC mothers, we found strong evidence for positive associations between polygenic scores for years of education and participation. This was observed consistently across all participation phenotypes (Figures 1 and 2, and Supplementary Figures S3- S5). Higher values of polygenic scores for height and agreeableness were also associated with higher participation across most participation phenotypes. There was also some evidence that higher polygenic scores for openness were associated with the mother completing more questionnaires about herself. In contrast, genetic risk scores for BMI, schizophrenia, ADHD, smoking initiation and depression were negatively associated with participation. Polygenic scores for neuroticism were associated with lower participation by the mothers (mother questionnaires, most recent clinic attendance and most recent questionnaire completion), but not with total participation. There was no clear evidence for an association between age at menarche and participation in the ALSPAC mothers.

**Figure 1.**
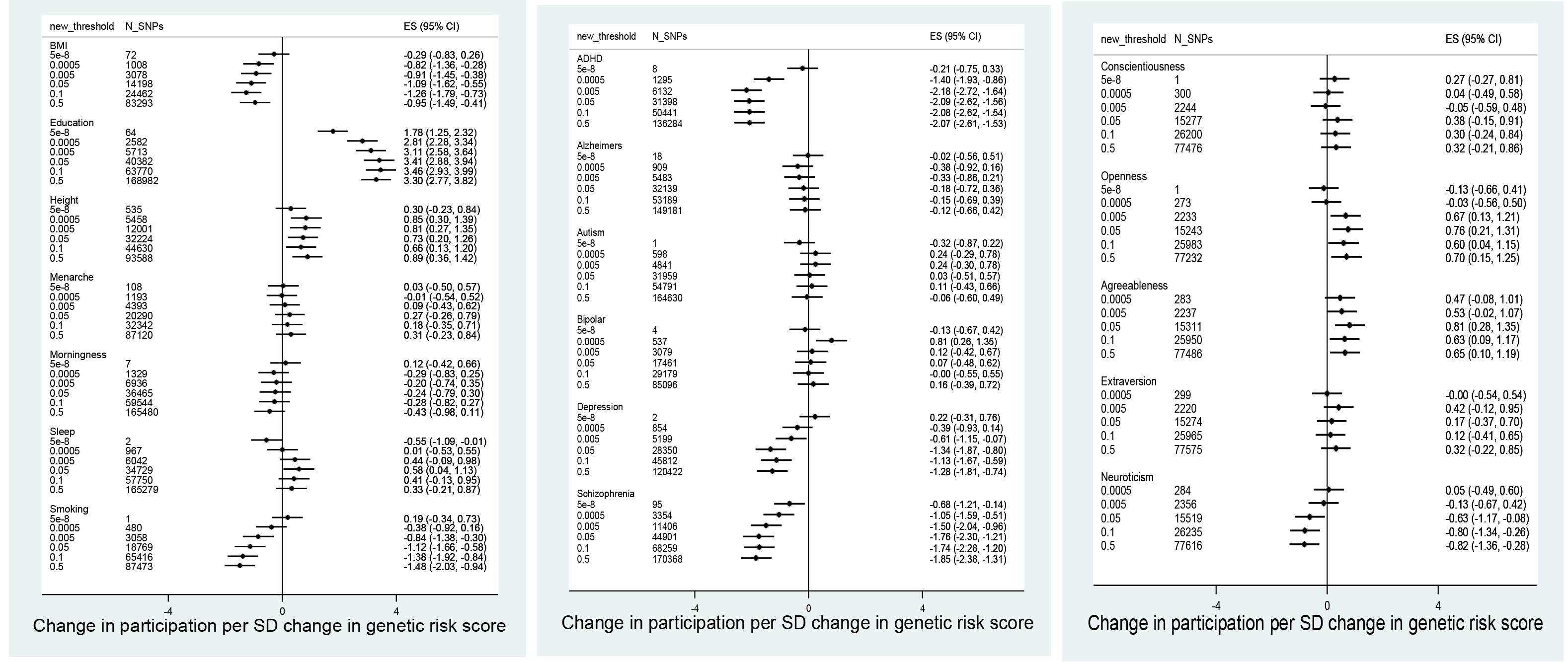
Association between polygenic scores in ALSPAC mothers and total participation score (N=7,468)

**Figure 2.**
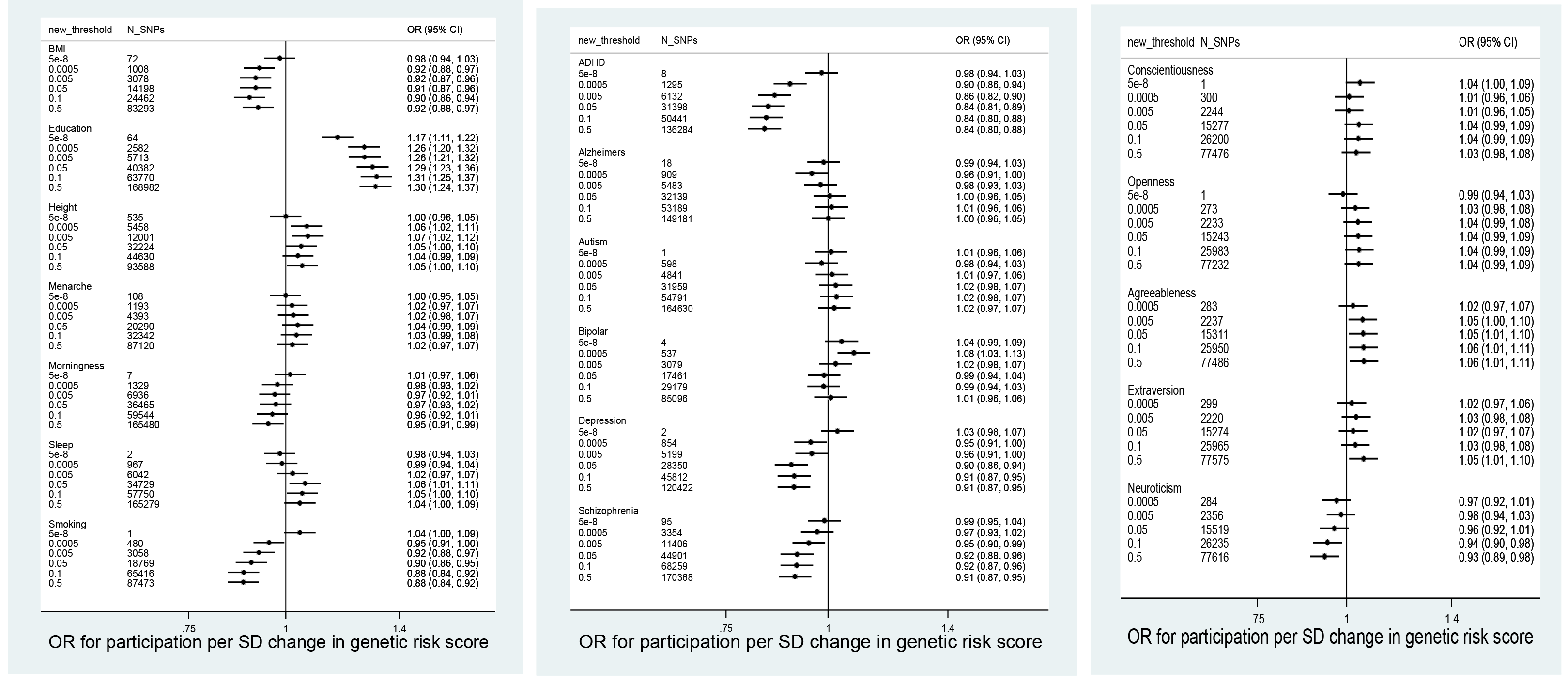
Association between polygenic scores in ALSPAC mothers and completion of most recent questionnaire (N=7,468)

Associations between polygenic scores and participation were similar for ALSPAC children (Figures 3 and 4 and supplementary figures S6-S9). Polygenic scores for education, height and agreeableness were positively associated with participation. Polygenic scores for smoking initiation, schizophrenia, ADHD and depression were negatively associated with participation. Genetic scores for age at menarche were positively associated with higher participation, with stronger evidence for the continuous measures than for last questionnaire and last clinic participation. In contrast to the ALSPAC mothers, there was little evidence for associations between polygenic scores for neuroticism or openness and participation.

**Figure 3.**
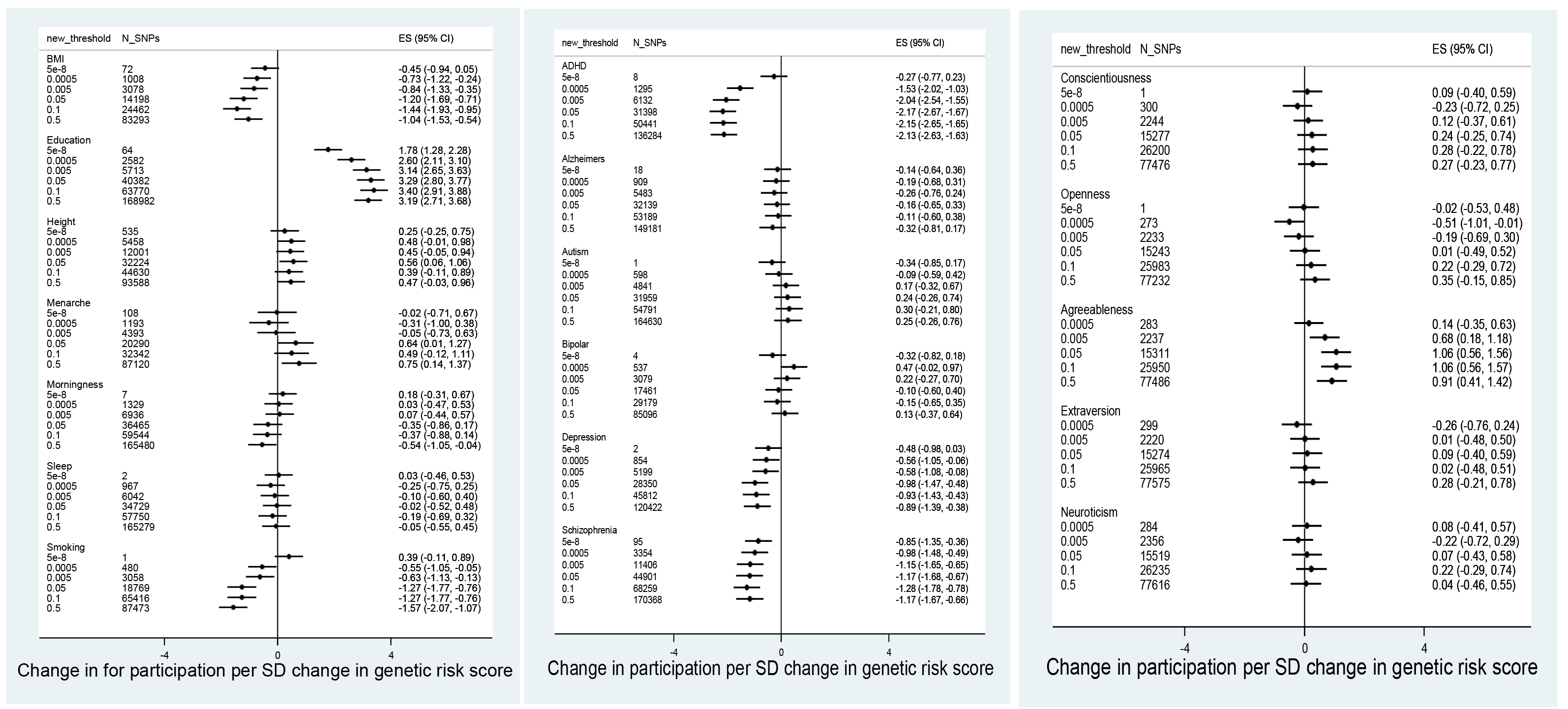
Association between polygenic scores in ALSPAC children and total participation score (N=7,508)

**Figure 4.**
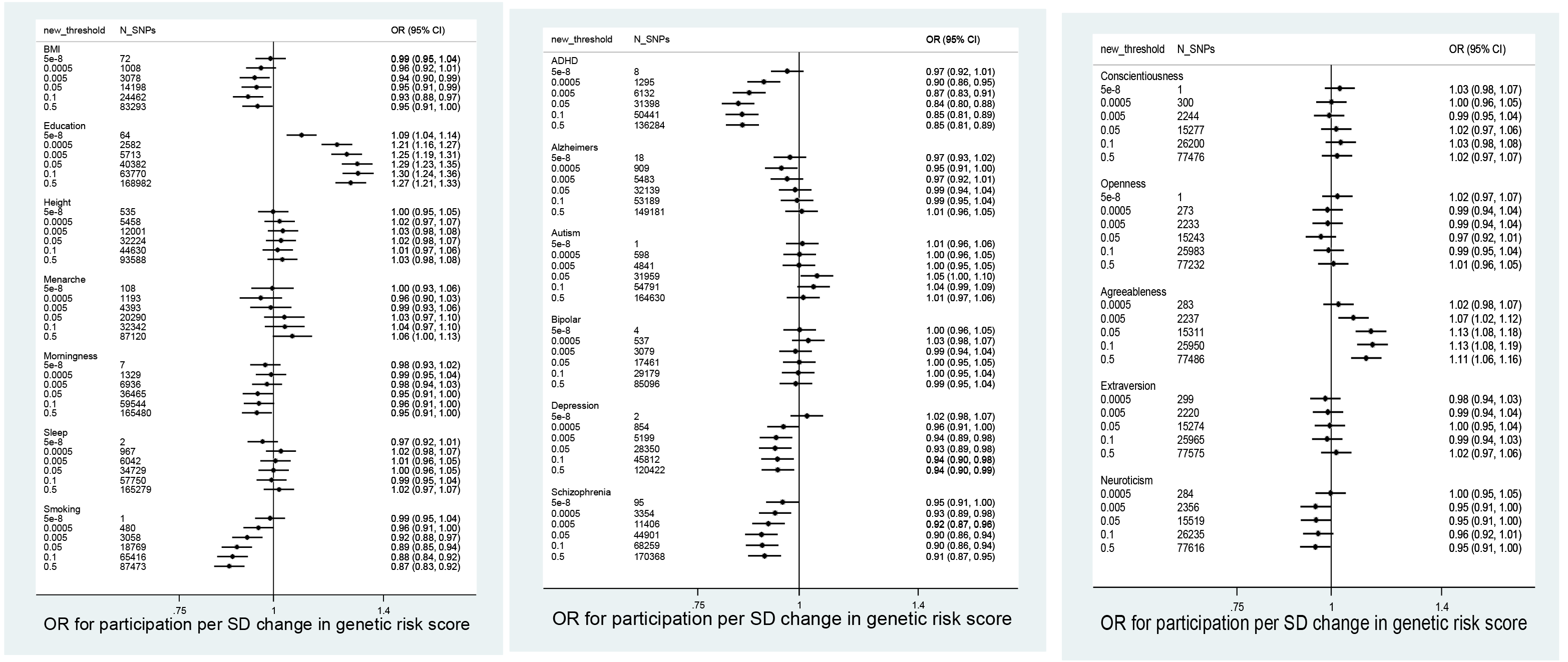
Association between polygenic scores in ALSPAC children and completion of most recent questionnaire (N=7,508)

We found no consistent evidence that polygenic scores for morningness (chronotype), sleep, bipolar disorder, autism, conscientiousness, extraversion or Alzheimer’s disease were associated with participation.

### Correlations between polygenic scores

The degree of correlation between polygenic scores for different traits at p<0.0005 and p<0.5 is shown in supplementary tables S7-S10. Correlations tended to be stronger for scores derived using the higher p-value thresholds.

### Investigating the impact of selection bias in ALSPAC

Figure 5 shows associations (in the largest sample with genome-wide data and in a sub-sample who attended the most recent clinic) between polygenic scores (constructed at the p<0.05 threshold) for BMI, smoking, education and schizophrenia and self-reported BMI and smoking. Associations between each polygenic score and smoking or BMI were in the same direction in both the full sample and the sub-sampleand in many cases of similar magnitude. However, associations between the polygenic score for education and being an ever smoker were substantially attenuated in the sub-sample (OR:0.95 per SD in polygenic score for smoking, 95% CI: 0.88, 1.02 compared to the full genetic sample (OR: 0.85, 95% CI: 0.81, 0.89). The association between the education polygenic score and BMI was also attenuated in the sub-sample compared to the full sample. In contrast, the association between the smoking polygenic score and BMI appeared stronger in the sub-sample compared to the full genetic sample, although the confidence intervals overlapped.

**Figure 5.**
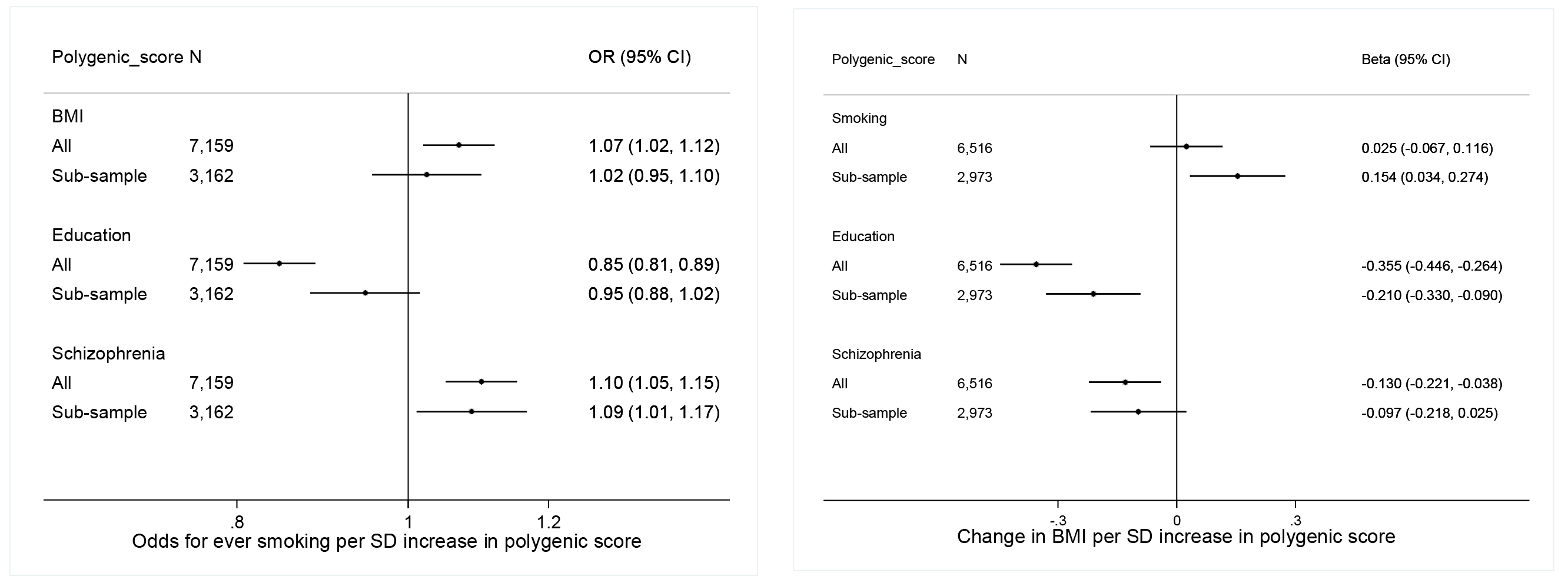
Association between genetic risk scores for BMI, smoking, education and schizophrenia and self-reported smoking and BMI, conditioned on attendance at the most recent clinic.

### Genome-wide association studies

Only one locus reached genome-wide significance with participation in the ALSPAC mothers and no genome-wide significant associations were found in the children (Figure 6, Supplementary Figure S10-S13). In the mothers, variants located in an intergenic region on chromosome 7: 51995163-52042976 were associated with total participation, total questionnaire and mother questionnaire (Figure 6, Supplementary Figure S10 and Tables S11-S13). Genome-wide hits were all in strong linkage disequilibrium (R^2^ > 0.8), indicating that this represents a single genetic signal. The SNP with the smallest p-value was rs10626545 for total (P=1.42x10^−9^) and total questionnaire (P=2.36e-9) and rs406001 for mother questionnaire (P=1.21x10^−8^). SNPs in this region reached genome-wide significance or close to genome-wide significance (P<7x10^−7^) with dichotomised total participation, total questionnaire and mother questionnaire (data not shown). However, the minor allele frequency of these variants was relatively low (0.012) and beta-coefficients large (beta for total participation for top SNP= 10.9), suggesting this association is driven by a few individuals.

**Figure 6.**
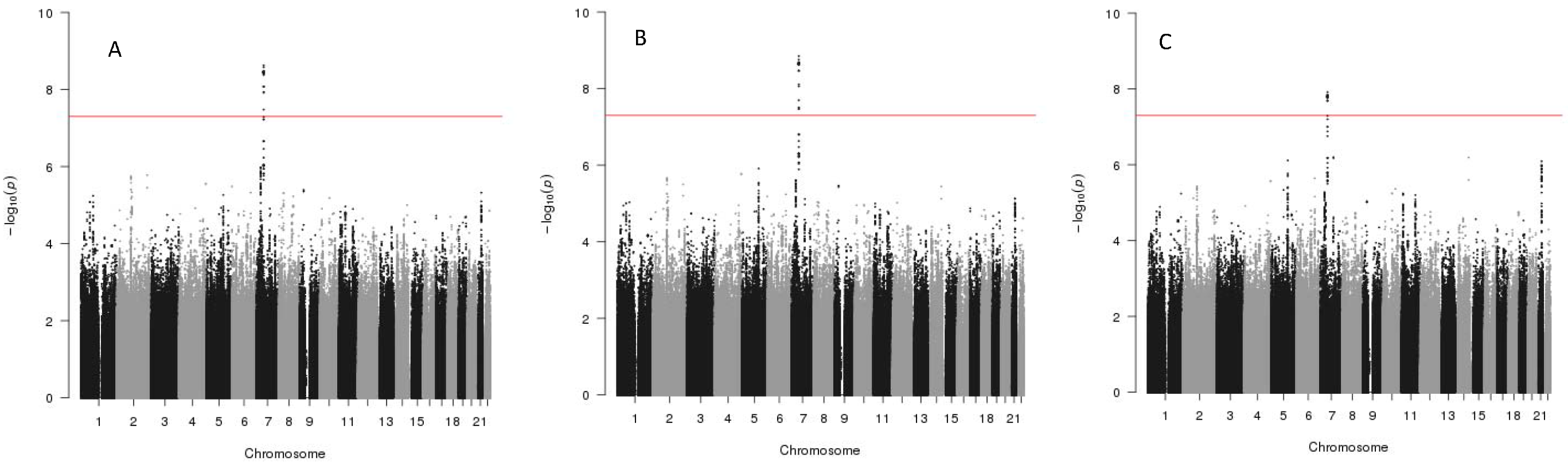
Manhattan plots for genome-wide analyses of total participation, total number of questionnaires completed and number of mother questionnaires in the ALSPAC mothers.

### SNP-based heritability

Estimates of heritability of participation phenotypes from SNPs included in the genome-wide analyses ranged from 20-27% for the mothers and 17-31% for the children (p-values all <0.001) (Supplementary Table S14).

## Discussion

Continued participation in the ALSPAC cohort is related to polygenic scores for a number of lifestyle factors, personal characteristics and health conditions, including level of education, BMI, height, smoking, agreeableness, openness, schizophrenia, ADHD and depression. We did not find robust evidence in genome-wide analyses that specific single genetic variants influence degree of participation in ALSPAC, though there was evidence of common genetic variants explaining a modest proportion of the variation in participation (up to 30%).

Our findings show that genetic variants which are related to specific phenotypes are also related to participation. Using a Mendelian randomisation framework this could imply that these phenotypes cause continued participation. For example, the polygenic risk score for education was the score most robustly associated with participation-implying that higher education causes greater continued participation in ALSPAC. This interpretation requires that the key assumptions of Mendelian randomisation are met (34), namely that: 1) the polygenic score is robustly associated with the trait of interest, 2) there are no confounders of the polygenic score-participation association, and 3) the genetic risk score only affects participation through the trait of interest. The third of these assumptions is more likely to be met as the threshold for polygenic score construction gets closer to genome-wide significance.

For polygenic scores created using higher p-value thresholds, polygenic scores are likely to be less specific for the trait of interest and more likely to be pleiotropic, influencing more than one trait. This is shown by the stronger correlations between risk scores for different traits created at high p-value thresholds than those created
using low p-value thresholds. We found traits for which genome-wide scores were not associated with participation, but scores at higher p-value thresholds were e.g., depression. This could be explained by low power in the original GWAS, meaning that truly associated SNPs are less likely to be included in a score constructed using a low significance threshold (35), or that effects on participation are acting through a trait which is only distally related to the GWAS trait used in score construction.

We also showed that it is possible to introduce bias into genetic analyses even when sample sizes are relatively modest. Therefore, we cannot assume that genetic-association studies, including GWAS, candidate gene studies and Mendelian randomisation are not biased by incomplete participation. We recommend that researchers consider how likely non-participation is as a potential source of bias when running genetic association studies and acknowledge this when reporting findings. The same implications hold for non-genetic studies – e.g. a study of the association between education levels and BMI in a selected sub-sample is likely to be biased by selection, since our genetic results show that both exposure and outcome cause participation (11).

For both genetic and non-genetic studies, there are potential methods to correct for this bias. For example, where there is some information about participants who have dropped out, it may be possible to apply inverse probability weighting (36). Where such data are not available, other approaches could be triangulated to examine likelihood of bias. Negative control exposures and/or outcomes can be used to see if associations between genetic variants and outcomes exist that are not biologically plausible and should only arise through selection bias (37). Similarly, where there is a well characterised association (replicated in a number of studies) of known magnitude between a genetic variant and an outcome, this can be used as a positive control. Finally, novel associations should be replicated in populations which have not undergone the same degree of selection.

We only found one locus associated with participation at genome-wide significance level. SNPs at this locus (e.g., rs406001) were identified in a previous GWAS of post-traumatic stress disorder (PTSD), but not replicated in the original GWAS (38). Furthermore, this SNP was only nominally associated with PTSD in a much larger GWAS (39). This, coupled with the low minor allele frequency of SNPs in the genome-wide significant locus in our GWAS suggests that this may be a chance finding, rather than an effect of PTSD on participation.

There are a number of limitations to this analysis. First, our analysis sample was restricted to just over half of the enrolled sample, due to availability of DNA samples for GWAS and exclusion criteria. Individuals in the analysis sample had higher participation rates than the full sample, meaning that associations between polygenic scores and participation are likely to be weaker than we would observe if we had full genetic data for the whole cohort. Second, our results may not be generalisable to studies with different selection criteria or specific cultural or contextual factors influencing participation. It is also possible that characteristics influencing participation will change over time and with age. Third, we have not attempted to disentangle the relative influence of maternal and offspring genetics on participation. It is likely that child participation is heavily influenced by maternal traits in childhood and this may continue into adolescence and adulthood. Finally, we have not explored all possible traits that might be associated with participation, since our analyses required access to GWAS summary statistics.

In conclusion, we demonstrate that polygenic scores related to a wide range of traits are associated with degree of participation in ALSPAC and that this may introduce bias into genetic and non-genetic analyses. This highlights the importance of considering selection bias in all studies, and the need for the development of statistical methods to account for this issue.

## Acknowledgements

We are extremely grateful to all the families who took part in this study, the midwives for their help in recruiting them, and the whole ALSPAC team, which includes interviewers, computer and laboratory technicians, clerical workers, research scientists, volunteers, managers, receptionists and nurses. The UK Medical Research Council and Wellcome (Grant ref: 102215/2/13/2) and the University of Bristol provide core support for ALSPAC. This publication is the work of the authors and will serve as guarantors for the contents of this paper. ALSPAC children were genotyped using the Illumina HumanHap550 quad chip genotyping platforms by Sample Logistics and Genotyping Facilities at Wellcome Sanger Institute and LabCorp (Laboratory Corportation of America) using support from 23andMe. AET and MRM are members of the UK Centre for Tobacco Control Studies, a UKCRC Public Health Research: Centre of Excellence. Funding from British Heart Foundation, Cancer Research UK, Economic and Social Research Council, Medical Research Council, and the National Institute for Health Research, under the auspices of the UK Clinical Research Collaboration, is gratefully acknowledged. This work was supported by the Medical Research Council (MC_UU_12013/1, MC_UU_12013/5, MC_UU_12013/6, MC_UU_12013/9) and the NIHR Biomedical Centre at the University Hospitals Bristol NHS Foundation Trust. The views expressed in this paper are those of the authors and not necessarily the MRC, Wellcome, NIHR or any other funders,

